# UNG-RPA interaction governs the choice between high-fidelity and mutagenic uracil repair

**DOI:** 10.1101/2024.04.30.591927

**Authors:** Yunxiang Mu, Zaowen Chen, Joshua B. Plummer, Monika A. Zelazowska, Qiwen Dong, Laurie T. Krug, Kevin M. McBride

## Abstract

Mammalian Uracil DNA glycosylase (UNG) removes uracils and initiates high-fidelity base excision repair to maintain genomic stability. During B cell development, activation-induced cytidine deaminase (AID) creates uracils that UNG processes in an error-prone fashion to accomplish immunoglobulin (Ig) somatic hypermutation (SHM) or class switch recombination (CSR). The mechanism that governs high-fidelity versus mutagenic uracil repair is not understood. The B cell tropic gammaherpesvirus (GHV) encodes a functional homolog of UNG that can process AID induced genomic uracils. GHVUNG does not support hypermutation, suggesting intrinsic properties of UNG influence repair outcome. Noting the structural divergence between the UNGs, we define the RPA interacting motif as the determinant of mutation outcome. UNG or RPA mutants unable to interact with each other, only support high-fidelity repair. In B cells, transversions at the Ig variable region are abated while CSR is supported. Thus UNG-RPA governs the generation of mutations and has implications for locus specific mutagenesis in B cells and deamination associated mutational signatures in cancer.

## Introduction

Uracils are common DNA base lesions that are created by misincorporation of dUTP and spontaneous or enzyme catalyzed deoxycytidine deamination^1^. The Apolipoprotein B mRNA editing catalytic polypeptide-like family (APOBEC family), including APOBEC3 members and activation-induced cytidine deaminase (AID), actively convert cytidine to uracil in ssDNA^2–4^. If left unrepaired, uracils derived from deamination are 100% mutagenic, causing C to T transition mutations when replicated over^5^. Mammalian uracil DNA glycosylase (UNG) is the primary uracil repair enzyme that recognizes and cleaves the uracil base to produce an abasic site (AP site)^6^. This triggers the high-fidelity base excision repair (BER) pathway to faithfully restore the genetic information. However, in B lymphocytes, UNG activity is co-opted for mutagenic repair to support immunoglobulin (Ig) gene diversification^7^. Furthermore, certain mutational signatures in cancer, such as Catalogue of Somatic Mutations in Cancer (COSMIC) signature Single Base Substitution 2 (SBS2) and SBS13, are derived from error-prone repair of cytidine deamination events catalyzed by APOBEC3 proteins^8–10^. The molecular mechanism for governing the choice between high-fidelity vs. mutagenic uracil repair is not understood^11^.

AID and UNG activities are crucial to antibody diversification. During the geminal center (GC) reaction, AID targets both the immunoglobulin (Ig) switch regions to initiate class switch recombination (CSR), and the Ig variable domains to achieve somatic hypermutation (SHM). During CSR, uracils within switch regions are processed by UNG and mismatch repair pathways to generate double strand breaks (DSBs) substrates for recombination. Ig isotype switching is accomplished through deletional recombination using a non-homologous end joining mechanism to change isotypes. During SHM, point mutations are introduced into the variable regions of the Ig genes that can alter antibody affinity. B cells that express antibodies with higher affinity are positively selected for during the process of affinity maturation. AID triggers SHM by introducing uracils in the variable region. The uracils are either processed by UNG or left unrepaired. Unrepaired uracil is replication competent and recognized as thymidine, resulting in a C to T transition. UNG excises the uracil base, leaving an AP site. AP sites lack genetic information and block high-fidelity DNA polymerases. Typically, the classic BER pathway repairs AP site in dsDNA efficiently, through sequential lesion processing by AP endonuclease 1 (APE1), DNA polymerase beta, and DNA ligases^12^. Alternatively, when an AP site occurs in ssDNA and base pairing information is absent, translesion DNA polymerases can bypass the AP sites. These low-fidelity DNA polymerases often incorporates the wrong nucleotides, resulting in C to T transitions, or C to G/A transversions^5,13^. Although the Ig variable locus is preferentially targeted by AID^14^, AID activity occurs at transcribed genes across the genome^15^. In contrast to mutagenic uracil repair at the Ig variable region, uracil repair at non-Ig genes preferentially maintains fidelity^15^.

UNG is widely represented in eukaryotes, prokaryotes as well as several large DNA viruses including herpesviruses (HVs). Gammaherpesviruses (GHVs) which include the oncogenic human herpesviruses (HHV), Epstein–Barr virus (EBV/HHV-4), Kaposi sarcoma associated herpesvirus (KSHV/ HHV-8) and the model murine Gammaherpesvirus 68 (MHV68/MHV-4) infect and maintain life-long latency in host B lymphocytes^16^. All GHVs studied to date encode a functional UNG^17–23^. In the murine model pathogen system, the viral UNG of MHV68 (MHV68UNG) promotes virus replication in the lungs^21,22^. MHV68 then migrates to secondary lymphoid organs where it latently expands in the GC and establishes life-long latency in the B cell compartment^22^. MHV68UNG has uracil glycosylase activity on ssDNA and dsDNA^23^ and compensates for CSR deficiency in UNG knockout (KO) B cells^21^. While GHVUNGs share structure and sequence identity with mammalian UNGs in their core catalytic domain, GHVUNGs diverge from mammalian counterparts in two domains: the N-terminal region, and an extension of the leucin loop structure in their DNA binding domain^20,23^. In GHVs, the unique leucine loop extension of UNG mediates a difference in DNA substrate choice. The GHVUNG leucine loop also imparts affinity to AP sites, a quality not found in mammalian UNGs^23^. During its life cycle, MHV68 inhabits GC cells. During the peak of latency, day 14-22 post infection, MHV68UNG protein is detected in a significant percentage of infected GC cells. Whether the MHV68UNG protein participates in or alters SHM or CSR is not known^24,25^.

Here we investigated the ability of MHV68UNG to impact cytidine deaminase mediated mutagenesis and found that it only supports high-fidelity repair. By exploiting the differential mutational outcome rendered by these UNGs, we identified the key characteristic of mammalian UNG that mediates error-prone uracil repair. We define the replication protein A (RPA) binding motif (RBM) of mammalian UNG and its interaction with RPA32 as critical molecular determinants that support error-prone repair.

## Results

### Differential uracil repair activity of mUNG and MHV68UNG

GHVUNG and murine UNG (mUNG) have divergent biochemical properties. GHVUNGs have affinity for AP site due to a unique leucine loop extension in their DNA binding domain^23^. MHV68UNG compensates for CSR in *ex vivo* cultured mUNG KO B cells to an extent that is similar to mUNG^21^. However, it is not known whether MHV68UNG supports hypermutation in a manner similar to mammalian UNG. To determine the effect of MHV68UNG on AID mediated somatic mutations, we utilized a cell line with a mutation target reporter gene. The fibroblast 3T3 cells (3T3-NTZ) has a Tet-off transcription-inducible GFP reporter with a stop codon integrated into its genome. This GFP gene undergoes mutation in an AID- and transcription-dependent manner; and the mutation frequencies and spectra are quantified by sequencing^26^. These cells have been utilized to determine factors that influence AID induced mutation frequency^27–29^. To directly compare uracil repair by mUNG and MHV68UNG, we transduced 3T3-NTZ cells with the indicated UNGs. The N-terminal FLAG tag facilitated detection of UNG protein, and FLAG-UNG expressing cells were selected with puromycin. The UNG expressing cells were then transduced with a retrovirus expressing AID and mCherry. The GFP reporter gene transcription was induced via removal of tetracycline. After 11 days, mCherry+ AID expressing cells were collected by FACS sorting and the GFP reporter gene was amplified by PCR, cloned and individual clones were evaluated for mutation by sequencing (Figure 1A). In an agreement with previous studies^26,27^, AID drove a high mutation frequency with most clones displaying multiple mutations (Figure 1B).

**Figure 1.**
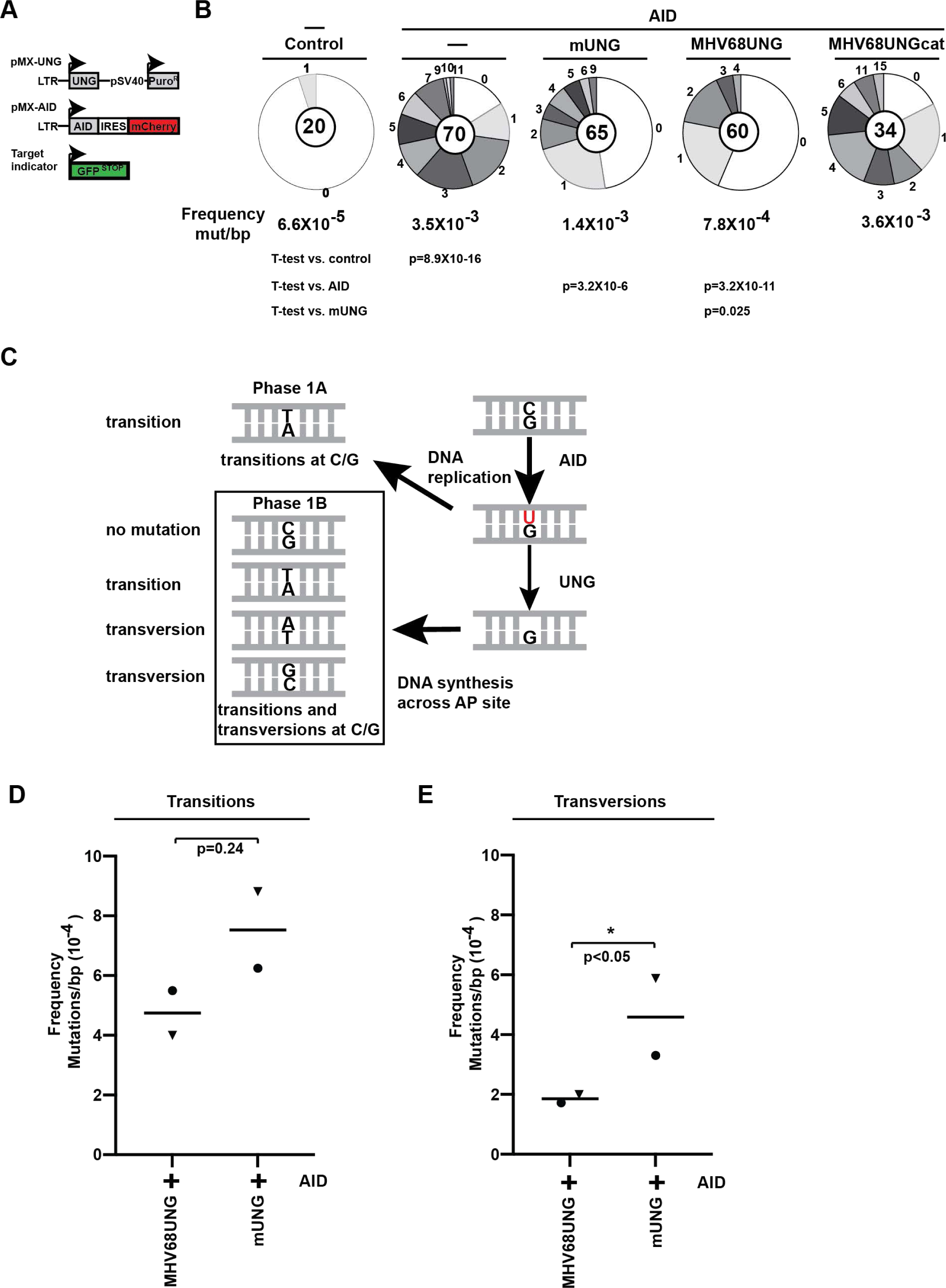
MHV68UNG suppresses hypermutation. (A) Diagram of mutation assay in 3T3-NTZ cells showing introduced expression retroviral plasmids and mutation target gene. (B) Number of mutations in mutation indicator gene cloned from fibroblasts expressing AID and indicated UNG. Segment size in pie chart is the proportion of clones with indicated number of mutations. Number of clones analyzed is indicated in center. Mutation frequency is displayed below pie chart (C) Model for how AID deamination and UNG activity give rise to transition or transversion mutations a deamination sites. Unrepaired uracil may be replicated over to give rise to transition mutations (Phase 1A). UNG cleaves uracil that is created by AID to produce an AP site, which can be bypassed by translesion polymerases to create transition, transversion or no mutation (Phase 1B). (D)Mutation frequency of transition at C/G (E) Mutation frequency of transversion at C/G. Two replicate experiments were performed (denoted by triangle and circle symbol). P-values (*, P < 0.05) were determined by a two-tailed t test assuming unequal variance.

Overexpressing mUNG is known to suppress mutation frequency^30,31^. However, we observed a significantly lower mutation frequency upon MHV68UNG expression compared to mUNG expression (MHV68UNG 7.8X10^-4^ mut/bp versus mUNG 1.4X10^-3^, p= 0.025) (Figure 1B). Expression of the catalytic mutant of MHV68UNG (D85N, H207L) led to comparable mutation frequency as the vector control. The observation that MHV68UNG and mUNG display different uracil repair activities on mammalian genomic DNA led us to analyze the spectra of AID induced mutations. C to T transition mutations arise from DNA replication over unrepaired U:G mismatches and these are suppressed by UNG uracil excision and high-fidelity BER repair (Figure 1C)^13^. Comparing C to T transitions frequency in the reporter gene we observed a modest, but insignificant decrease upon the expression of MHV68UNG compared to mUNG (Figure 1D). In contrast, MHV68UNG caused a significant drop in transversion mutation frequency greater than 2-fold (Figure 1E). Transversion mutations (C to A or G) reflect UNG processing followed by error-prone repair (Figure 1C)^13^. Analysis of uracil-DNA glycosylase (UDGase) activity in cell lysates demonstrated that the lower mutational frequency of MHV68UNG was not due to greater activity (Figure S1). mUNG expressing cells displayed higher UDGase activity on uracil containing substrates. Altogether, these results suggest that MHV68UNG processed genomic uracil in a high-fidelity manner more efficiently than mUNG.

### Mammalian UNG and GHVUNG differ at the N-terminus and leucine loop extension

Mammalian and GHV UNGs have structural and sequence conservation within their core catalytic domain. Significant divergence is noted in two areas: the N-terminal domain, and a leucine loop extension motif within their DNA binding domain (Figure 2). The GHVUNGs leucine loop extension is longer than mammalian UNGs with a unique sequence identity. We recently reported that the leucine loop extension confers an affinity to AP sites, a characteristic that the mUNG lacks^23^. In the nuclear isoform of mammalian UNG (UNG2), N-terminal motifs mediate interaction with partner proteins such as proliferating cell nuclear antigen (PCNA), via the PCNA-interacting protein-box (PIP-box) and Replication protein A 32 (RPA32), via the RPA binding motif (RBM).

**Figure 2.**
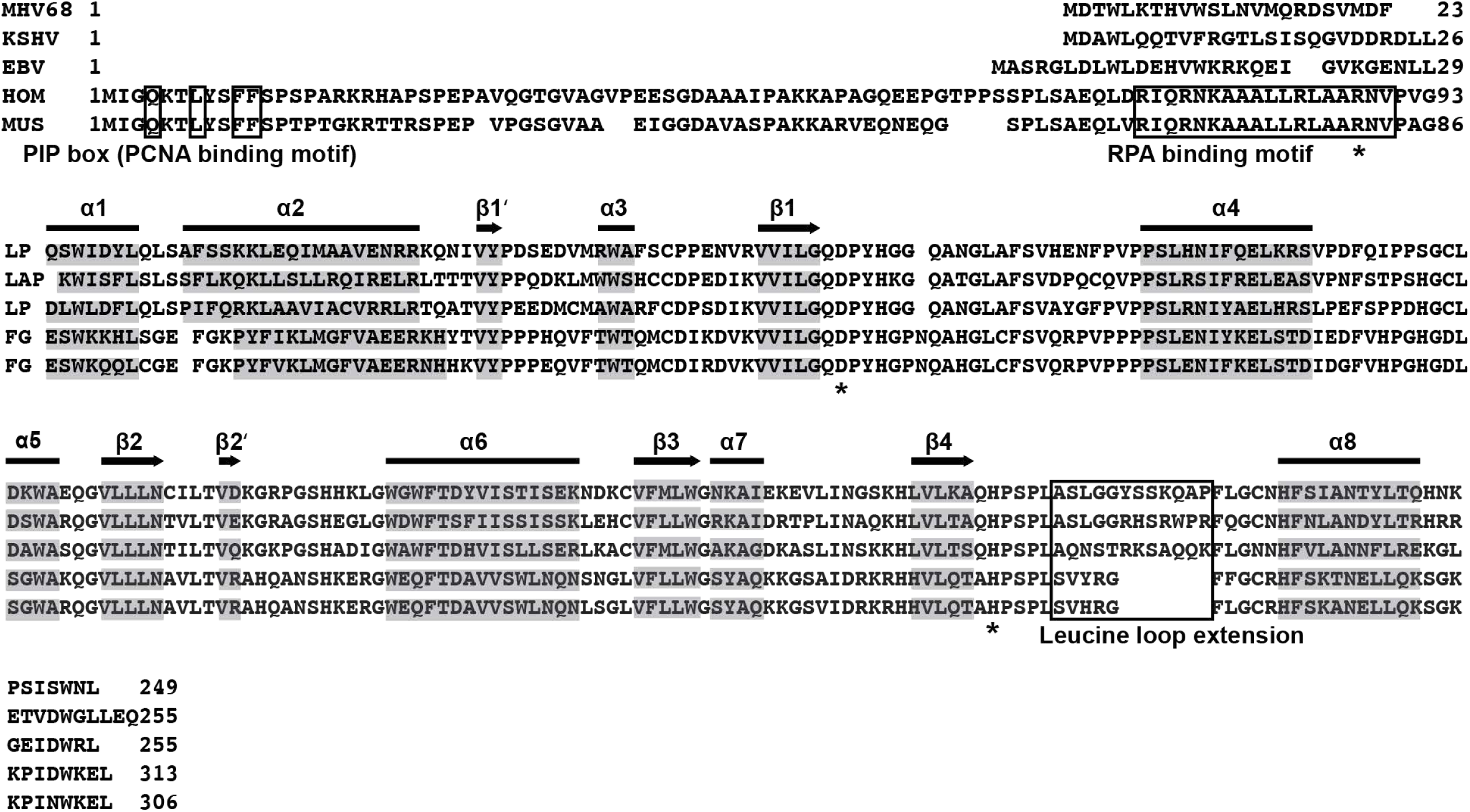
Alignment of gHV UNGs and mammalian UNGs. Alignment of murine gammaherpesvirus (MHV68), Human Kaposi sarcoma herpesvirus (KSHV), Epstein-Barr virus (EBV), Human (HOM) and mouse (MUS) UNGs. Denotation of secondary structural elements, alpha helix (horizontal bar) and beta sheet (arrow) are based on PDB structures of KSHV (PDB ID: 5NNU), EBV (PDB ID: 2J8X), and human UNG (PDB ID: 1UGH). Sequence of these elements are shaded. Catalytic site aspartic acid (MHV68UNG D85) and histidine MHV68UNG H207) residues are labeled with asterisk. R81 in mouse UNG and R88 in human UNG are denoted with asterisk. PCNA binding motif, RPA binding motif (RBM), and leucine loop extension motif are boxed.

These motifs are absent in the N-terminus of GHVUNGs (Figure 2).

### Leucine loop extension of UNG does not affect UNG repair outcome

To determine whether the N-terminal or leucine loop extension structures contribute to differential mutation outcomes, we constructed chimeric UNG mutants with swapped domains^23^. Utilizing the NTZ reporter assay we compared the frequency of mutation in cells expressing mUNG, MHV68UNG or mutants with their leucine loops swapped (Ls) (Figure 3A). No significant difference in the overall mutation frequency (Figure S2A) or either the transition or the transversion mutation frequency was observed between the wild-type MHV68UNG or mUNG and their corresponding Ls mutants (Figure 3B, 3C).

**Figure 3.**
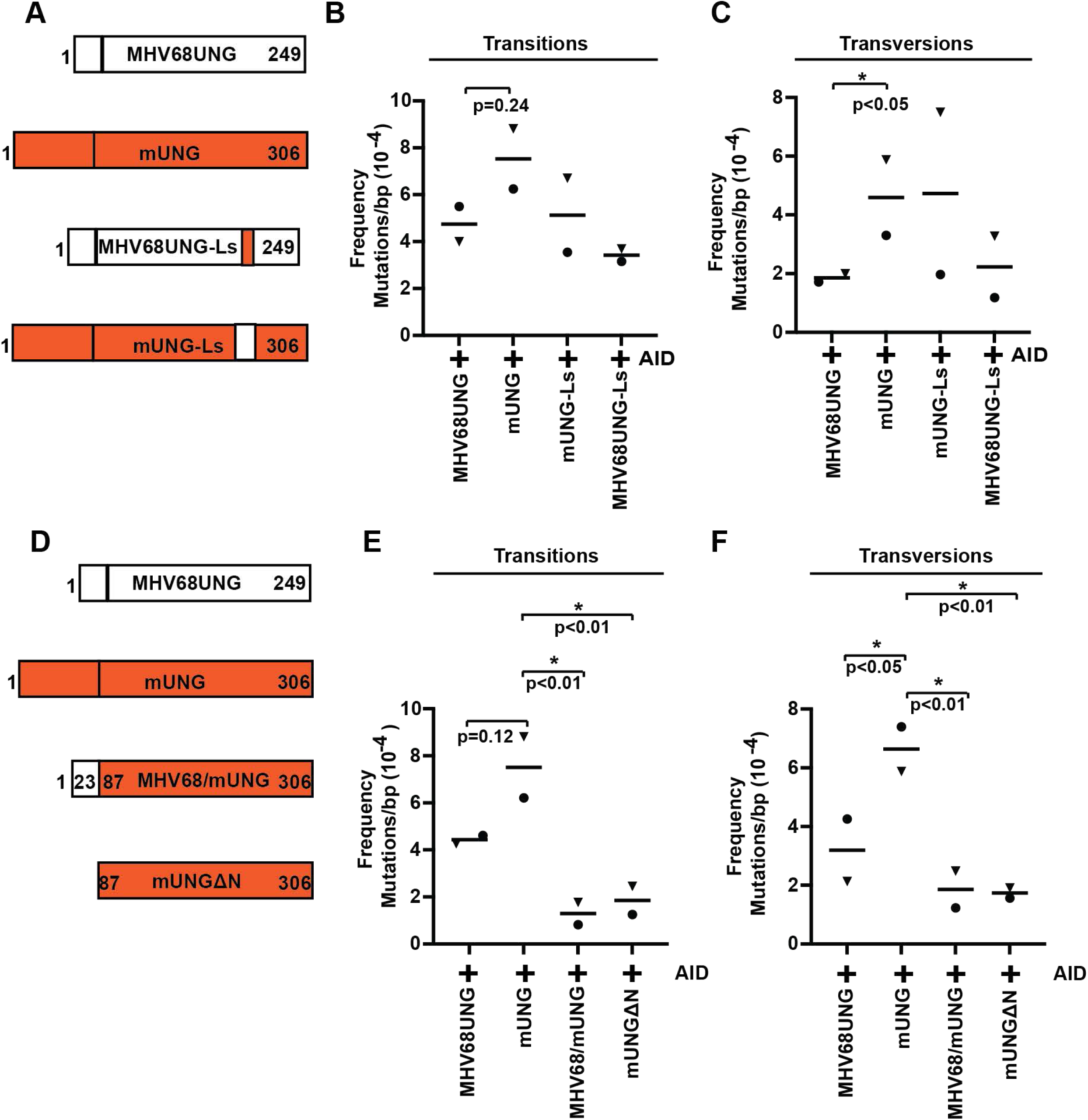
The N-terminus of mUNG is the determinant of transversion mutations. (A) Diagram of UNG chimeras at the leucine loop extension with white representing MHV68UNG sequence and orange denoting mUNG (B) Transition and (C) Transversion profiles of mUNG Leucine loop swap mutants. (D) Diagram of UNG N-terminus mutants (chimera and truncation mutant) (E) Transition and (F) Transversion profiles of mUNG N-terminal mutants. Two replicate experiments were performed (denoted by triangle and circle symbol). P-values (*, P < 0.05 or P < 0.01) were determined by a two-tailed t test assuming unequal variance.

### N-terminus of mUNG contributes to error-prone uracil processing

We next examined whether the N-terminus of mUNG contributes to mutagenic repair. Cells expressing a mUNG mutant with the N-terminus deleted (mUNGLlN) or mutants with swapped N-terminal regions were analyzed (Figure 3D). Cells expressing mUNG with the N-terminus of MHV68UNG (MHV68/mUNG) or mUNGLlN displayed a drop in overall mutation frequency compared to mUNG (Figure S2B). The frequencies of transition mutations of the mUNGLlN (1.9X10^-4^ ts/bp) and MHV68/mUNG (1.3X10^-4^ ts/bp) were significantly decreased compared to mUNG (7.5 X10^-4^ ts/bp) (p<0.01) (Figure 3E). Notably, the frequencies of transversion mutations of the N-terminal mutants, mUNGΔN (1.7X10^-4^ tv/bp) and MHV68/mUNG (1.9X10^-4^ tv/bp) were also significantly decreased compared to mUNG (6.6 X10^-4^ tv/bp) (p<0.01) (Figure 3F). Importantly, this difference in mutational frequency was not due to differences in enzymatic activity, since our previous analysis of catalytic activity demonstrated that mUNG and mUNGΔN had comparable catalytic activity on both ssDNA and dsDNA uracil containing substrates and MHV68/mUNG had slightly lower activity^23^. This result suggests that the N-terminal domain of UNG plays a major role in dictating error-prone repair outcome.

### Mutagenic uracil repair by mUNG relies on RPA32 interaction

The N-terminus of mammalian UNG contains a PIP-box and an RBM (Figure 2). To determine if either motif contributes to mutagenic uracil repair, we utilized a UNG mutant with altered residues in the PIP-box motif known to disrupt the UNG-PCNA interaction, (mUNG F10A/F11A)^32^ and a RPA binding mutant (mUNG R81C)^33^. Mutation analysis in the NTZ reporter cell line revealed similar mutation frequency in cells expressing mUNG versus mUNG F10A/F11A (Figure S2C). In contrast, the mUNG R81C expression led to a significant decrease in mutation frequency (4.2X10^-4^ mut/bp) compared to mUNG (8.6 X10^-4^ mut/bp, p=0.024) (Figure S2C). In comparison to mUNG, both transition and transversion mutation frequency were significantly decreased (Figure 4A) (mUNG 6.3 X10^-4^, R81C 1.8 X 10^-4^ ts/bp, p<0.01; mUNG 5.2 X10^-4^, R81C 3.0 X 10^-4^ tv/bp, p<0.05). To confirm that the R81C mutation conferred a reduction of interaction with RPA, a co-immunoprecipitation (co-IP) analysis was performed. The UNG proteins expressed in the reporter cell lines contained a FLAG epitope tag. Anti-FLAG immunoprecipitation revealed a reduced interaction between mUNG R81C and endogenous murine RPA32 while MHV68UNG did not interact with murine RPA32 (Figure 4B). To determine if the UNG R81C mutant was also deficient in supporting CSR we examined its ability to recapitulate CSR in B cells from UNG^-/-^ mice. Control vector, mUNG, mUNG R81C or a catalytic mutant of mUNG (mUNGcat - D147N, H270L) were introduced retrovirally into UNG^-/-^ B cells. Cells were cultured *in vitro* in the presence of LPS and IL4 to induce CSR to IgG1. While non-transduced, vector control and mUNGcat displayed background levels of CSR, mUNG and mUNG R81C induced IgG1 switching to similar levels (Figure 4C and 4D). The specific activity of mUNG R81C was similar to wild-type mUNG on uracil containing substrates in the context of ssDNA or dsDNA (Figure 4E). These results indicate that UNG deficient in RPA interaction can process AID induced uracils but displays impaired mutagenic repair.

**Figure 4.**
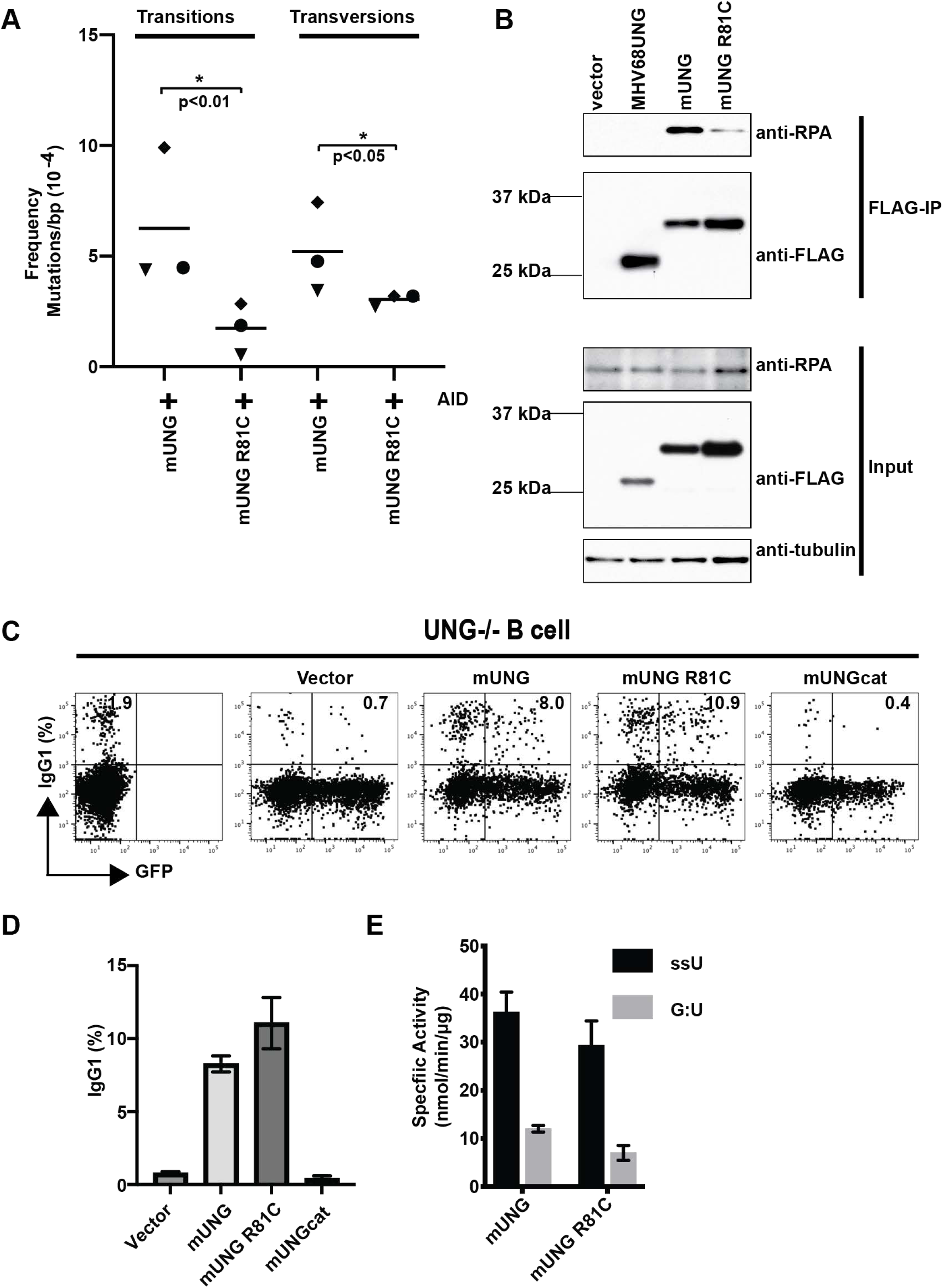
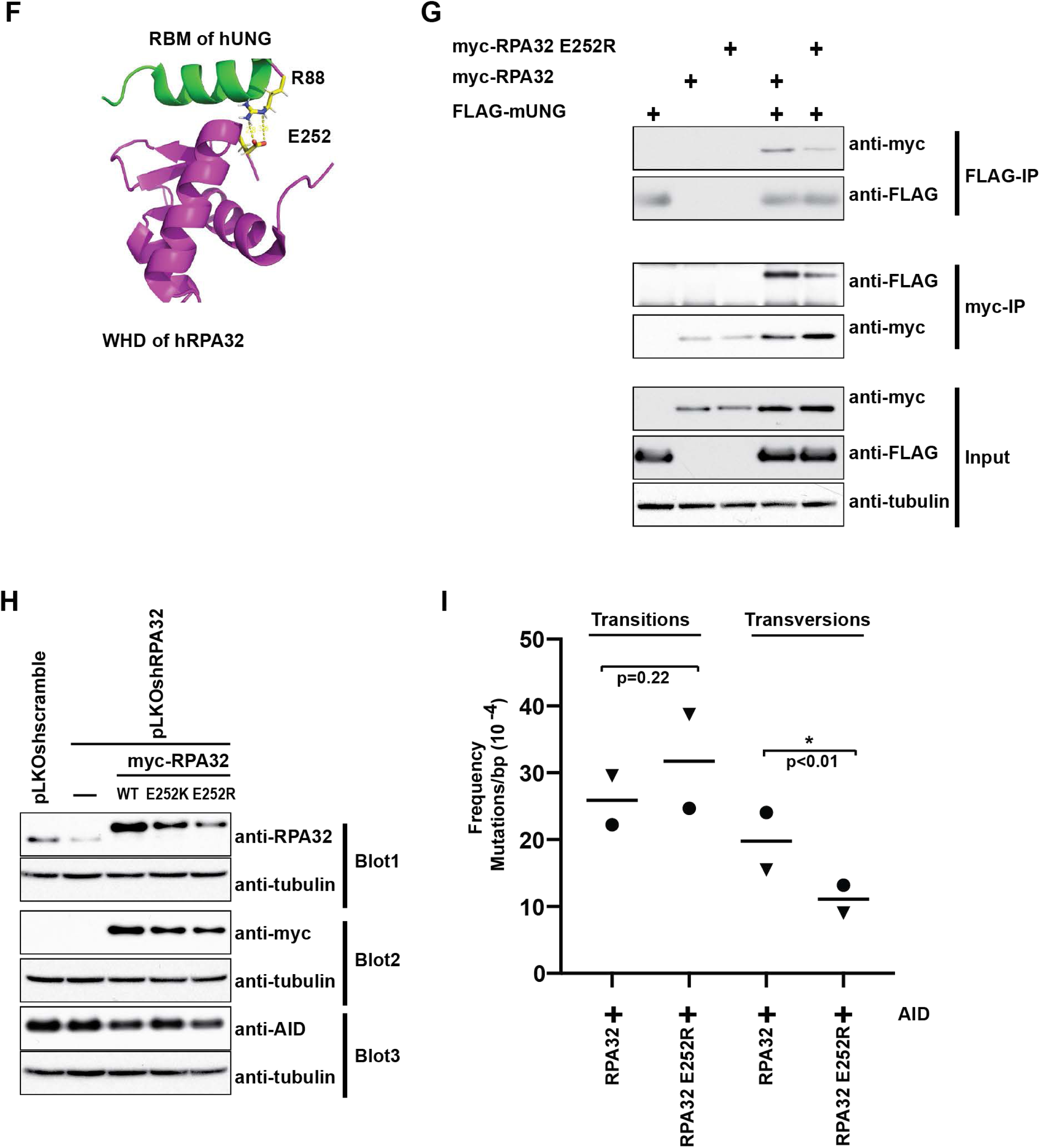
RPA-UNG interaction determines transversion mutations. (A) Transition and transversion mutation frequencies in mutation assay of cells expressing mUNG and mUNG R81C. Three replicate experiments were performed (denoted by triangle, diamond and circle symbol). (B) Anti-Flag co-IP of RPA in cells expressing Flag-MHV68UNG, -mUNG or -mUNG R81C. Immunoblots displaying input with tubulin input control and co-IP are displayed. Representative of n=3 experiments. (C) Representative flow cytometry plot of CSR to IgG1 in UNG^−/−^ splenocytes transduced with control vector with -IRES-GFP or expressing -mUNG, -mUNG R81C, or -mUNGcat (catalytic mutant). The percentage of infected (GFP+) cells expressing IgG1 is indicated on the plot. (D) Summary graph of percentage of UNG^−/−^, GFP+ cells that underwent CSR to IgG1 3 days after transduction from n=3 experiments. (E) Specific activity of recombinant mUNG and R81C on indicated uracil containing substrates in oligo breakage assay. (F) NMR structure of peptide corresponding to the RBM of hUNG in complex with the WHD of human RPA32 (PDB:1DPU). The hUNG-R88 and RPA-E252 that interact through an electrostatic interaction are visualized. (G) Anti-FLAG and anti-myc tag co-IP from expressing cell lysates. Transfection of epitope-tagged mUNG and mRPA32 are indicated. Antibody used from IP and immunoblot of input lysates are indicated. Representative of n=2 experiments. (H) Immunoblots of NTZ cell lysates from lentivirus control or shRNA knockdown of RPA. RPA replacement was via transduction with retrovirus expressing indicated myc-mRPA32 WT or mutant. Cells used for immunoblot were from the same pool of cells that were also analyzed for mutation profile. Three immunoblots were performed on the same set of cell lysates. Loading control (anti-tubulin) for each is displayed. (I) Transition and transversion mutation frequency profiles from cells expressing wild-type and RPA32-E252R. Two replicate experiments were performed (denoted by triangle and circle symbol). P-values (*, P < 0.05) were determined by a two-tailed t test assuming unequal variance

### RPA32 mutant with decreased UNG interaction supports lower transversion mutation

To ensure that the observed phenotype of mUNG R81C was due to the reduced RPA interaction, we sought to define the contribution of RPA to UNG mediated error-prone repair. We characterized an RPA mutant (E252R) deficient in UNG binding. The rationale for mutating glutamate 252 is based on the NMR structure of the winged-helix domain (WHD) of human RPA32 (hRPA32) in complex with the RBM of human UNG (hUNG) (Figure 4F)^34^. UNG interacts with RPA32 through an induced alpha helix of RBM, which is stabilized by an electrostatic interaction between hUNG R88 (equivalent to mUNG R81) and hRPA32 E252. We reasoned if glutamate 252 was replaced by a positively charged amino acid, such as arginine, it would repel R81 on mUNG disrupting the electrostatic bond with it. To determine if the E252R mutation disrupted the interaction with UNG, we performed reciprocal co-IP analysis. Reduced interaction was observed in cell lysates expressing RPA E252R when either Myc or FLAG antibodies were used for IP (Figure 4G).

We assessed the effect of RPA E252R on mutation. RPA is essential for cell survival, so we utilized a knockdown-replacement strategy by targeting a 3’ UTR of RPA32 mRNA with shRNA and reintroducing myc-tagged RPA32 in the reporter cell line^35^. In those cells, only the transduced N-terminal myc-tagged mRPA32 was detected as a slower migrating band, with no endogenous RPA32 detected (Figure 4H). Reporter cell mutation analysis revealed that RPA32 and RPA32 E252R supported a similar transition mutation frequency. In contrast, transversion mutation frequency in cells expressing E252R (11.1 X 10^-4^ tv/bp) was half that of cells expressing the wild-type RPA (19.8 X10^-4^ tv/bp, p<0.01) (Figure 4I). The overall higher mutation frequencies in this experiment compared to those with overexpression of UNG mutants reflect the endogenous level of UNG expression in RPA32 knockdown-replacement cell lines.

### UNG-RPA interaction determines transversion mutations in B lymphocytes

We utilized the hypermutating human Raji B cell to analyze the influence of RPA-UNG interaction on SHM at the Ig variable region. A CRISPR-HR mediated knock-in strategy was employed to create Raji clones where UNG contained a homozygous mutation for arginine 88 to cysteine, the equivalent of mUNG R81C (Figure 5A, 5B, S3A, S3B).

**Figure 5.**
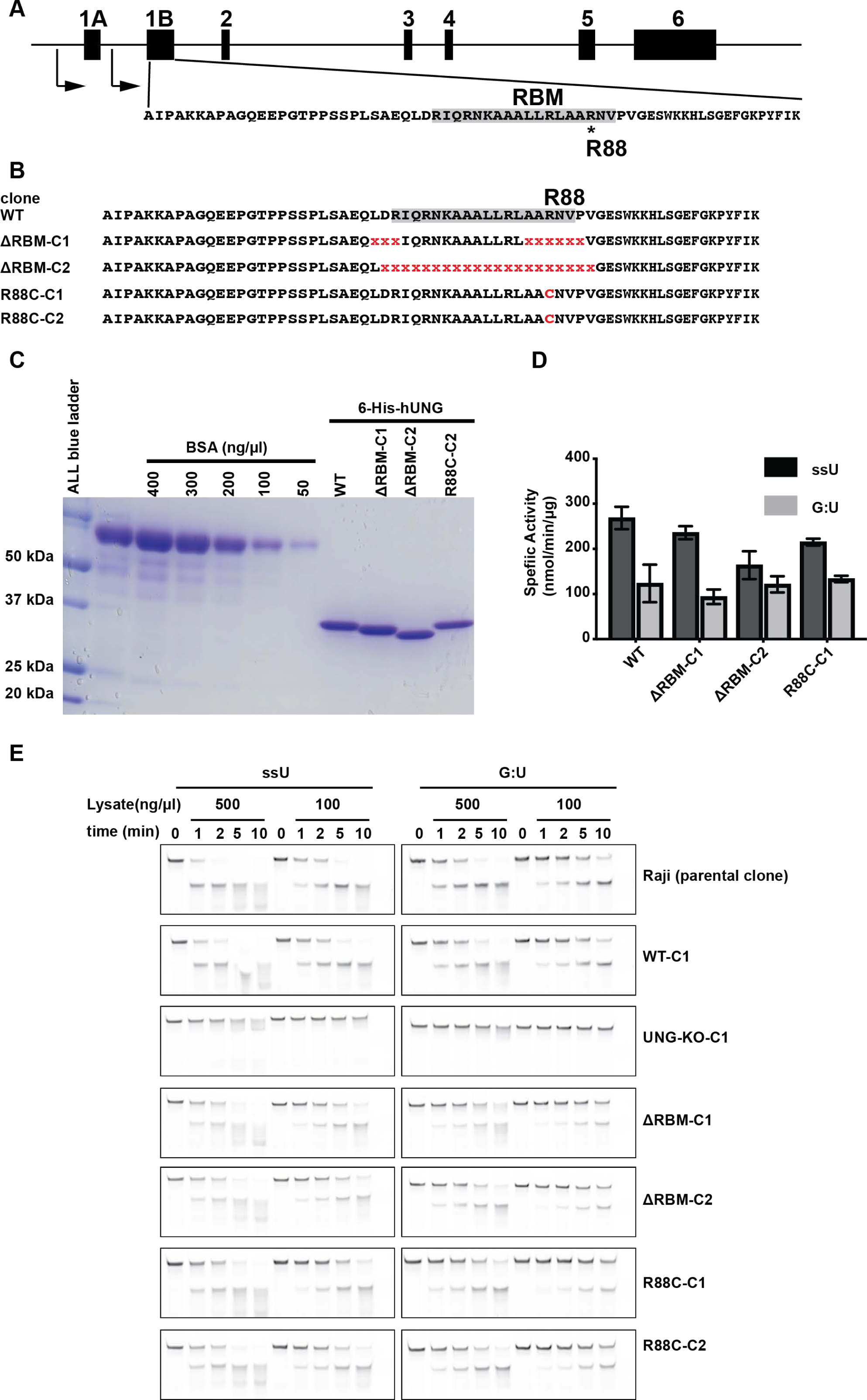
The RBM deletion mutant and R88C of hUNG do not affect enzymatic activity. (A) Schematic of the human UNG locus exons (black boxes) and RBM (shaded) within exon 1B is displayed. (B) Alignment sequences from clones following editing strategy in Raji cells. Deletions and R88C mutation indicated in red. (C) Coomassie staining of SDS-PAGE resolving his-tag purified recombinant hUNG mutants along with serial diluted BSA standard (D) UDGase activity of recombinant hUNG mutants on indicated uracil substrates from oligo breakage analysis. (E) Oligo breakage assay measuring UDGase activity in cell lysates from indicated Raji clones. Displayed is the conversion of ssU or G:U 36-mer uracil substrate to a 14-mer after indicated incubation time with lystate amount.

Clones that had in-frame deletions that removed part (ΔRBM-C1) or the entire UNG RBM (ΔRBM-C2) were also generated. The UNG R88C or UNG ΔRBM-Cs were recombinantly produced and purified (Figure 5C), and confirmed to be catalytically active on uracil containing ssDNA (ssU) and dsDNA (G:U) substrates in an oligo breakage assay similar to wild-type hUNG (Figure 5D). To ensure the Raji clones with mutant UNGs displayed similar UDGase activity, lysates were made and analyzed for activity on ssU and G:U substrates (Figure 5E). While a Raji UNG-KO clone displayed no UDGase activity, the mutants displayed similar activity as wild-type.

Raji cells were induced to undergo higher levels of Ig variable region hypermutation by culturing with a combination of anti-CD21, anti-CD19 antibodies and crosslinking of the surface BCR^36,37^. After 11 days of stimulation, mutation frequency and spectra of the expressed VDJ (IGHV3-21, IGHJ4) segment were determined via PCR amplification, cloning into plasmid and sanger sequencing of individual plasmid clones. VDJ mutation frequency was measured on 2-3 single clones in each group with 3 experimental replicates for each single clones. UNG knockout clone displayed a high frequency of C to T transitions (Figure 6A) compared to wild-type and UNG mutants. This was an expected consequence of AID deaminase activity in the absence of UNG repair activity^7^. Wild-type, R88C and ΔRBM clones had comparable transition mutation frequencies. This result suggested that the UNG mutants were processing AID induced uracils comparable to wild-type UNG. A robust frequency of transversion mutations were detected in all Raji clones with wild-type UNG (Figure 6B). The UNG-KO clone displayed background levels of transversion mutation (0.4X 10^-4^ tv/bp) near the expected frequency introduced by PCR error-rate. Both ΔRBM (0.5X 10^-4^ tv/bp, p<0.001) and R88C (1.2X 10^-4^ tv/bp, p<0.05) displayed significantly lower transversion frequency than wildtype hUNG (4.0X 10^-4^ tv/bp. The ΔRBM transversion frequency was similar to the UNG knockout frequency (UNG-KO) and near the expected PCR error-background frequency.

**Figure 6.**
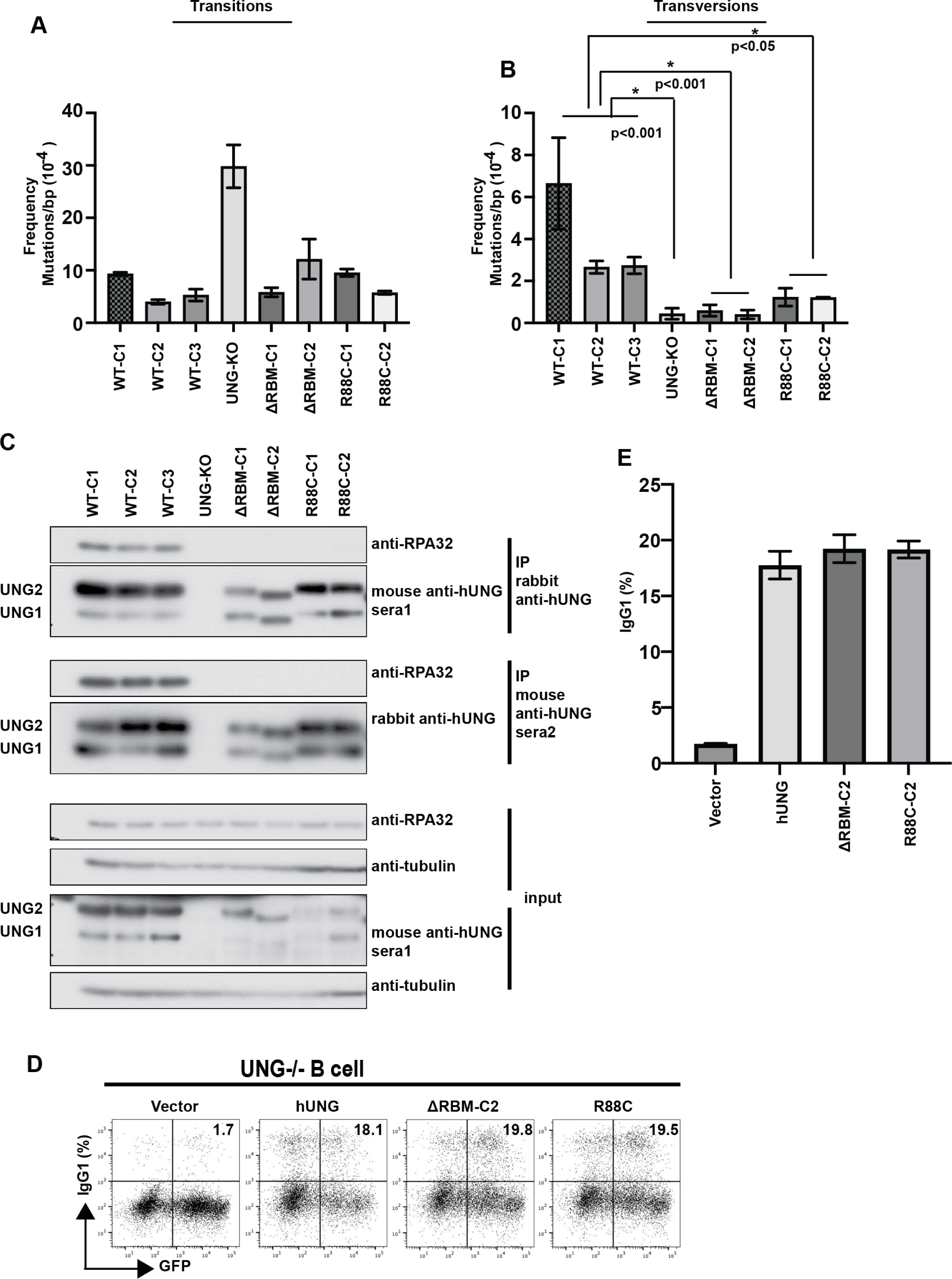
RPA-UNG interaction determines transversion mutations in B lymphocytes. (A) Transition and (B) transversion frequency profile from Raji clones with the indicated homozygous hUNG mutants. (C) Co-IP and immunoblot of Raji lysates. Two separate IP performed with indicated anti-hUNG antibodies are displayed. The migration of UNG1 (mitochondria isoform) and UNG2 (nuclear isoform) are indicated. (D) Representative flow cytometry plot of CSR to IgG1 from cultured UNG^−/−^ splenocytes transduced with retrovirus (GFP) control or expressing hUNG, ΔRBM, or R88C. The percentage of infected (GFP+) cells expressing IgG1 is indicated on the plot. (E) Summary graph of percentage of GFP+ cells that underwent CSR to IgG1 3 days after retroviral infection from n=3 experiments. Bars represent the means ± the standard deviations. P-values (*, P < 0.05 or P < 0.001) were determined by a two-tailed t test assuming unequal variance.

To measure the UNG-RPA interaction in the Raji cells, co-IP experiments were performed. We utilized antibodies that clearly recognized both hUNG1/2 isoforms in immunoblot (Figure S4) and were capable of pulling down the UNG-RPA complex in a co-IP. Two anti-hUNG polyclonal antibodies (commercial rabbit and in-house mouse sera) were found to be capable of co-precipitating the UNG-RPA complex (Figure 6C). hRPA32 was detected interacting with wild-type hUNG but not ΔRBM or R88C. Therefore, disruption of the RPA-UNG interaction results in ablation of transversion but not transition mutations of the Ig variable region in Raji cells. The ΔRBM expressing cells displayed only background transversion mutation frequency, while the R88C displayed frequency above background levels. When mUNG R81C was overexpressed, a decreased but detectable interaction with mRPA32 was revealed (Figure 4B). To determine if this was also the case for hUNG R88C, we cloned the WT, ΔRBM, R88C hUNG from cDNA of the Raji samples and expressed them as FLAG-tagged proteins in 293T cell line. IP with anti-FLAG beads revealed that the hRPA32 interaction with R88C was decreased but the interaction was ablated with hUNGΔRBM and mUNG-ΔRBM (Figure S5). This suggests that R88C retains a low degree of hRPA32 binding and is consistent with intermediate level of transversion mutations in R88C expressing clones.

### CSR is unaffected by loss of RPA interaction

We next determine if the UNG mutants support CSR upon expression in UNG KO mouse splenic B cells. Upon stimulation of CSR to IgG1 by culturing with LPS and IL4 *in vitro*, we observed a similar percentage of IgG1 positive cells in the samples expressing the WT UNG and UNG mutants (ΔRBM-C2 and R88C-C2) (Figure 6D,6E). These results suggest transversion mutations require UNG-RPA interaction (Figure 1C) while CSR does not.

## Discussion

Antibody maturation requires mutagenic repair to diversify AID created uracils in Ig genes^38^. Two mechanistically diverse repair outcomes are indispensable for SHM at the variable region and CSR at the switch region of Ig genes^39^. SHM entails UNG creating an AP site followed by error-prone repair (Figure 1C)^7^. Rev1 and other error-prone DNA polymerases are believed to bypass AP sites creating C to G or C to A transversion mutations at the C/G sites of AID targets^40–42^. Further processing of U:G mismatches by non-canonical mismatch repair factors^43–49^. engaged with PCNA K164 mono-ubiquitination^50–52^ and DNA polymerase eta^53–57^ permits error-prone repair at A/T sites flanking uracil. CSR requires DSBs. Single-stranded breaks are created by AP endonuclease processing of AP sites in CSR^58,59^. DSBs are thought to arise when breaks are in close proximity on opposite strands.

The catalytic domain of GHV UNGs and mammalian UNGs is conserved, the divergent leucine loop extension confers AP site interaction and DNA substrate specificity to GHV UNGs^23^. Built on *in vitro* experimental evidence, a “pass the baton” model for a faithful BER pathway has been proposed to efficiently repair damaged bases and protect genome stability^60,61^. In this model, DNA substrates with lesions or repair intermediates are channeled or handed over from the upstream player to the downstream player to ensure repair fidelity. Therefore, high affinity to AP site by lesion specific DNA glycosylases could block error-prone repair factor access and enable hand-over to high-fidelity pathway factors. Based on our initial finding that GHVUNG had AP site affinity, we hypothesized that this quality contributed to high-fidelity repair^23^. However, this is not the case. The leucine loop swap mutant of MHV68UNG (MHV68UNG-Ls) does not have AP site affinity^23^, but displays similar transversion frequency as WT. While *in vitro* analysis of human glycosylases and BER dynamics suggested that AP-site affinity could contribute to BER efficiency and repair outcome, our cellular mutation model did not support this for UNG^60^. Instead, we found that the N-terminus of mammalian UNG governed mutagenic uracil repair^62^.

We surmise that interactions with co-factors are the basis of mutagenic uracil repair. Since mammalian UNGs have the PIP-box and the RBM that are not present in GHVUNGs, we investigated their role. PIP-box mutant did not impact mutation frequency while RBM mutants affected transversion mutation. RBM mutants compensate CSR in UNG-KO B cells to a similar level as WT mammalian UNG. This contrasts with a recent report, using the murine CH12F3 B cell line, implicating a role for UNG-RPA interaction in CSR^63^. In that study, authors analyzed a single clone that expressed mUNG with a two amino acid in-frame deletion. That mutant ablated RPA interaction and displayed lower frequency of CSR to IgA. Several caveats could contribute to the different observations regarding the role of UNG-RPA in CSR. A clonal effect of this finding cannot be excluded and there may be a differential contribution of RPA to IgG1 versus IgA CSR. Our CSR compensation assay overexpressed UNG, which could also mask a CSR deficiency. Analysis of a mouse knock-in model could provide additional clarity.

A mechanism that diverts uracil repair to error-prone outcome in SHM has long been elusive^11^. In this study, using B cell and non-B cell lines, we demonstrate that UNG-RPA interaction is the determinant for conversion of uracil to a transversion mutation. A mouse mutation reporter cell line overexpressing AID and UNG with the R81C point mutant demonstrated the role of the UNG RBM. By constructing the corresponding mutation in the UNG binding motif of RPA32 (E252R) we corroborated that transversion mutations require the RPA-UNG interaction. Using the human Raji B cell hypermutation model, we provide further evidence for RPA-UNG interaction driving transversion mutations. Disrupting the UNG RBM through either in-frame deletions or R88C (equivalent to murine R81C) point mutation, abated interaction with endogenous RPA, and ablated transversion mutations, thus demonstrating that mutagenic repair of uracil in VDJ exon of the Ig variable domain is dependent on UNG-RPA interaction.

Transversion mutations arise when AP sites are channeled away from high fidelity BER to translesion bypass. In general, APE1 incises AP sites in dsDNA, triggering high fidelity BER. This process uses the complementary strand as a template for repair fidelity. APE1 has low activity on AP sites in the context of ssDNA, and it is known that RPA inhibits APE1 activity *in vitro*^64,65^. In the context of ssDNA, the RPA WHD domain facilitates UNG access and removal of uracil in ssDNA^66^. We speculate that the UNG-RPA interaction further remodels RPA, liberating the AP site and facilitating its access to translesion (TLS) polymerases such as Rev1^67^. Such a model is supported by RAD52, a homologous recombination factor, and RPA binding on ssDNA. RAD52 and UNG share a conserved RBM that interacts with the RPA32 WHD. Through single molecule analysis, it has been shown that the RAD52 RBM interacts with the WHD of RPA^68^. When ssDNA is coated by RPA, RAD52 binding causes RPA remodeling, allowing access to the ssDNA by other downstream repair factors^69^. RPA also facilitates UNG activity in dsDNA at the ssDNA-dsDNA junction^70^. If this occurs at a replication fork, an AP site will be converted into a ssDNA context through continued fork migration. This would be expected to leave the AP site susceptible to TLS bypass and conversion into mutation.

GC B cells express factors that promote error-prone repair in both a global and locus specific manner^14,15^. The level of UNG expression is a factor, lower levels promote CSR and SHM. B cells express FAM72A, which mediates UNG proteasomal degradation. In the absence of FAM72A, UNG protein levels increase, attenuating CSR, transition and transversion mutation frequency^71–73^. In our 3T3-NTZ cell line model, despite high expression levels, we observed distinct differences between mUNG and MHV68UNG. Using the hypermutating Raji cell line, which expresses physiologically relevant levels of hUNG, we defined the RBM of UNG to be the determinant of mutagenic repair. In GC cells, SHM is targeted to the Ig variable region^14^. While AID activity is focused on the Ig locus, AID deamination also occurs at non-Ig genes. However, the majority of these non-Ig AID targets are protected from mutation through high-fidelity repair^15,74,75^. How error-prone repair is controlled in a locus-specific manner has not been delineated^14^ but could involve regulating RPA-UNG interaction. Factors such as cell cycle stage^76^ and sequence context^31,75,77^ may also contribute.

Error-prone uracil repair has implications for cancer evolution and drug resistance development^78–80^. COSMIC signature SBS13 is one relevant case^8,9^. It is thought to be generated by translesion bypass after UNG processes the product of APOBEC deamination^8,9^. The SBS13 signature creates resistance mutation in some non-small cell lung cancers during targeted therapy^79^. Mutations that impart tyrosine kinase inhibitor resistance are found in EGFR (C797S: T<u>G</u>C to T<u>C</u>C)^79^ and ALK (S1206C:T<u>C</u>C to T<u>G</u>C; L1122V: <u>C</u>TG to <u>G</u>TG; G1269A: G<u>G</u>A to G<u>C</u>A)^79^. Therefore, understanding how error-prone repair is triggered can impact intervention strategies.

## Methods

### Cell lines, expression plasmids, and Lentiviral and retroviral production

BOSC cells and 3T3-NTZ cells were cultured in DMEM (Sigma) supplemented with 10% FBS (Corning, catalog # 35-010-CV) and Penicillin (10 units/ml)-Streptomycin (10 μg/ml) in CO_2_ incubator with 5% CO_2_ supply at 37 °C. Raji cells were cultured in RPMI1640 (Sigma, catalog # R8758-500ml) supplemented with 10% FBS, Penicillin-Streptomycin (Sigma, catalog # P4333-100ML), 1 mM Sodium pyruvate (Sigma, catalog # S8636-100ML), 10 mM HEPES (Sigma, catalog # H0887-100ML), and 50 μM 2-mercaptoethanol (Sigma, catalog # M3148-100ML). Cell lines were authenticated by STR DNA fingerprint and confirmed mycoplasma free (The University of Texas MD Anderson Cancer Center Cytogenetic and Cell Authentication Core).

Retroviral expression was via the pMX vector with expressed proteins inserted into the EcoR1 and Xho1 sites of the multiple cloning site (MCS). Untagged AID, UNGs with N-term Flag tag, murine RPA32 with N-term myc tag was expressed from pMX-IRES-GFP or derivatives (IRES-mCherry, -RFP670 or delta GFP). Primers are listed in Table S1. pMX vector, retrovirus production and infection has been described previously for overexpressing proteins in B cell and non-B cells ^27^. To make retrovirus that overexpresses UNG or RPA in 3T3-NTZ cells, 5 μg pMX-UNG-deltaGFP or pMX-RPA32-RFP670 was co-transfected together with 5 μg pCL-Eco into BOSC cells at 75% confluence in 10 cm plate using jetPrime transfection reagent (Polyplus, catalog # 101000046). Virus supernatant was collected by filtering cell culture media through 0.45 μm syringe filter (Millex® 33mm PVDF .45μm Sterile RUO, MilliporeSigma, catalog # SLHVR33RS) at 48 hours or 72 hours after co-transfection. Oligos with 5’ phosphate that target murine RPA32 3’ UTR were annealed with its complementary strand and ligated into pLKO vector at Age1 and EcoR1 sites. For lentivirus production, 5 μg of pLKO-ShRNA (scrambled or targeting RPA32 3’ UTR) and 3.75 μg of psPAX2, and 1.25 μg of pMD2.G were co-transfected into BOSC at 90% confluence in 10 cm plate. Media were replaced with fresh media 6 hours after co-transfection. Virus supernatant was collected by filtering cell culture media through 0.45 μm syringe filter (Millex® 33mm PVDF .45um Sterile RUO, MilliporeSigma, catalog# SLHVR33RS) at 48 hours or 72 hours after co-transfection. 3T3-NTZ were transduced with Retrovirus or lentivirus in the presence of polybrene (10 μg/ml) and 8 mM HEPES. For primary splenocyte transduction, B cells (2X10^5^ cells/ml) were transduced with retrovirus as described for 3T3 cells but underwent centrifugation at 90 min at 1150g at 29°C and incubation for 6 hours, before replacement with fresh media.

### Cell culture, retroviral infections and SHM assay

3T3-NTZ cells were transduced with control or Flag-tagged UNG expressing retrovirus, selected with 2 μg/ml puromycin (sigma, catalog # P8833-25MG) for 48 hours and then transduced with pMX-AID-IRES-mCherry retrovirus. After overnight AID retroviral transduction, regular DMEM were replaced with Tet-free DMEM to induce transcription of the genomic GFP mutation target indicator gene. Eleven days later mCherry+ cells were FACS sorted. Mutation analysis of a 939 bp region of the GFP indicator gene has been described ^26,27^. In brief, the target region of the GFP gene was amplified with Pfu-Turbo (Stratagene), cloned into TOPO-PCR4 (Thermo Fisher Scientific, catalog #r K458001) and individual colonies were sequenced. SHMtool was used for mutation spectrum analysis ^81^. Student T-test was applied for statistical analysis.

To test the role of RPA in hypermutation, 3T3-NTZ cells were transduced with Lentiviral pLKO-ShRPA32 targeting 3’UTR of murine RPA32, selected with puromycin for 48 hours. Retrovirus expressing RPA32-IRES-RFP670 (WT or E252R) was transduced into RPA32 knockdown 3T3-NTZ cells. These were then transduced with AID-mCherry retrovirus. RFP670 and mCherry double positive cells were FACS sorted.

### Mice and CSR assay

Ung^−/−^ mice (Endres et al., 2004) on C57BL6/J background were utilized as splenocyte sources. Animals were bred and housed at the accredited University of Texas MD Anderson Cancer Center animal facility. All work was approved by the Institutional Animal Care and Use Committee in accordance with the National Institutes of Health guidelines for use of live animals. Naïve B cells were purified from splenocytes upon CD43 depletion as previously indicated ^27,82^. CSR was induced by culture of purified naïve B cells in the presence of LPS (Sigma, catalog # L4130-100mg) and IL4 (R&D systems, catalog # 404-ML). UNG^-/-^ B cells were transduced with retrovirus expressing indicated UNG mutants. 72 hours after transduction, B cells were stained with anti-Mouse IgG1-APC (BD Pharmingen, catalog # 560089) and analyzed with a BD Fortessa cytometer. Data were analyzed with Flowjo (10.9.0)

### CrispR-cas9-HR repair mediated UNG knock-in, single cell sorting and repair junction analysis by direct PCR and sequencing

We sorted single B cells from Burkitt lymphoma Raji cell line (ATCC, catalog# CCL-86) and performed cell line authentication and mycoplasma check. A single clone was expanded and utilized for knock-in generation. That clone was sent to Synthego Corporation (Redwood, Ca) which generated the knock-in UNG mutants, RBM in-frame deletions and R88C point mutant from it. Pools of Raji knock-in cells were received from Synthego. We sorted single clones in 96-well plates, expanded the clones via culture and analyzed for state of the UNG gene in each clone. Direct PCR strategy was applied to each single clone without genomic DNA extraction using Phire Tissue Direct PCR Master Mix (Thermo Fisher Scientific, Catalog#: F170S) under the following cycling conditions: 98°C for 5 min, then 40 cycles of 98°C for 15 s, 56°C for 15 s, 72°C for 1 min, and 72°C for 7 min. Alternatively direct PCR can be performed using cells as DNA template with Platinum™ Direct PCR Universal Master Mix (Thermo Fisher Scientific, Catalog#: A44647500) under the following cycling conditions: 94°C for 2 min, then 35 cycles of 94°C for 15 s, 60°C for 15 s, 68°C for 1 min, and 68°C for 2 min. Primers used were SynthegoF: 5′-GGGCTCTTACTGTCCGCTTT-3′, and SynthegoR: 5′-ACTTGGCCGGCTACACTAAC-3′. PCR products were directly sequenced without purification using primer SynthegoF: 5′-GGGCTCTTACTGTCCGCTTT-3′.

For clones containing candidate homozygous RBM deletions or R88C mutations, the entire open reading frame (ORF) coding the cDNA human UNG2 (nuclear isoform) was confirmed. The cDNAs encoding hUNG L1RBM and R88C were also cloned into the pMX or pET-DUET-1 vectors. 1X10^7^ cells from each single clone were used for RNA extraction with RNeasy Plus Mini Kit (Qiagen, catalog#, 74134). cDNA was synthesized using SuperScript™ III Reverse Transcriptase (Thermo Fisher Scientific, Catalog#: 18080-093) and primer hUNG3 5′-GATTCTGGAAAGTTGCAAAACATC-3′.

cDNA synthesis was followed by nested PCR with 1^st^ round PCR by primer set of hUNG5 5′-TGACCGCCACAGCCACA-3′, and hUNG3 5′-GATTCTGGAAAGTTGCAAAACATC-3′ and 2^nd^ round PCR by either primer set of Duet1hUNG2EcoR1FP 5′-ggatccgaattcgATGATCGGCCAGAAGACGC-3′, and Duet1hUNG2Sal1RP 5′-ggaagcgtcgacTCACAGCTCCTTCCAGTCA-3′ for cloning into pET-DUET-1 or primer set of FLAG-hUNG 5′-ATGGACTACAAGGACGACGATGACAAGGGAGGAATCGGCCAGAAGACGC-3′ Xho1STOPhUNGR 5′-ggccgcctcgagTCACAGCTCCTTCCAGTCA-3′ followed by additional round of PCR with primer set of ECOR1KOZFLAG 5′-CGGATCCGAATTCCACCATGGACTACAAGGACGAC-3′ and Xho1STOPhUNGR 5′-ggccgcctcgagTCACAGCTCCTTCCAGTCA-3′ for cloning into pMX vector.

### Mutation analysis in Raji cells

To induce high levels of hypermutation, 2X10^6^ Raji cells in log phase growth were incubated with 4 μg/ml biotin labeled goat anti-human IgM (Thermo Fisher Scientific, catalog #, #31778), 1 μg/ml mouse anti-human CD19 (SouthernBiotech, catalog #, 9340-01), 5 μg/ml mouse anti-human CD21(BD Biosciences, Catalog #, 555421) in 500 ml of basic RPMI medium at 4 °C for 20min. The cells were washed twice with basic RPMI and incubated with 15 μl of streptavidin magnetic beads (Thermo Fisher Scientific, catalog #, #88816) in 500 μl of basic RPMI medium at 4 °C for 20 min with rotation. Cells were resuspended in 4 ml B cell medium and cultured for 11 days with log phase growth. To harvest, Raji cells were pelleted and washed twice with PBS. Genomic DNA was prepared by PCI extraction twice (Phenol:Chloroform:Isoamyl Alcohol 25:24:1 Saturated with 10 mM Tris, pH 8.0, 1 mM EDTA, Sigma, catalog # P2069-100ML) followed by isopropanol (Sigma, catalog # I9516-500ML) precipitation and 70% ethanol wash. Air dried genomic DNA were resuspended in milli-Q water and quantified with DeNovix Ds-11 FX Spectrophotometer/Fluorometer. The VDJ region (IGHV3-21*06-j4) expressed by Raji was amplified with PfuTurbo (Agilent, catalog # 600252) under the following cycling conditions: 98°C for 1 min, then 30 cycles of 98°C for 10 s, 63°C for 30 s, 68°C for 1 min, and 68°C for 7 min. Each group was amplified in triplicates, polished by Taq polymerase (Sigma, catalog # D1806-1.5KU) at 72°C for 20 min, agarose gel purified, pooled, and cloned into pCR4-TOPO (Invitrogen), and individual colonies were sequenced. Statistical significance was determined by two-tailed Student’s t test assuming unequal variance. Primers used were hVH-Raji-F: 5′-GGAGGCCTGGTCAAGCCG-3′, hVH-Raji-R: 5′-TTACACGGAGAGACATTGCGAGGA-3′. For mutational analysis background mutations were noted and not included in the mutation frequency.

### Protein purification from *E. Coli* and UDGase activity assays

Evaluation of UDGase activity of recombinant UNG has been previously described ^23,83^. UNGs were cloned into EcoR1 and Sal1 sites in pET-Duet-1 vector. BL21 (DE3) *E. Coli* were transformed for recombinant protein expression. Expression of recombinant 6His-tagged murine and human UNG mutants was induced with 0.2 mM IPTG at 18°C overnight. Cells were lysed in lysis buffer (20 mM Tris, pH 7.5 and 150 mM NaCl plus protease inhibitors), sonicated, clarified, and incubated with Cobalt resin (Pierce) overnight. Following wash with lysis buffer,1% triton and 5 mM Imidazole in lysis buffer, protein was eluted with 150 mM imidazole, dialyzed into 20 mM Tris pH 7.5, 25 mM NaCl, 1 mM EDTA, 1 mM DTT, 10% glycerol and frozen at -80°C. recombinant UNGs were quantified with BSA standard using SDS-PAGE and image J analysis. Glycosylase activity was evaluated using a 36-nt dsDNA oligo with the substrate oligo containing a fluorescein-dT label and internal uracil (AGAAAAGGGGAAAGUAAAGAGGAAAGGTGAGGAGGT) ^23,83^ 333 nM substrate was incubated with UNG in 50 mM Potassium Acetate, 20 mM Tris-acetate, 10 mM Magnesium Acetate, 1 mM DTT pH 7.9 for indicated time, quenched with PCI and purified with phase lock gel, treated with 0.1 M NaOH at 95C for 5 minutes and separated by 16% Urea-PAGE. Specific activity was calculated from three replicates experiment and plotted with Prism.

### Production of polyclonal sera against human UNG

Purified 6-his tagged human UNG-RBM mutant ((flRBM-C2) was used to raise polyclonal antibodies against human UNG in C57/BL6 mice. The mouse was immunized with 50 μg of recombinant hUNG-flRBM-C2 in Imject™ Alum Adjuvant (Thermo Scientific, Catalogue #. 77161). The primary immunization was through IP (Peritoneal injection). At two-week intervals, the mouse received boost of 20 mg protein Imject™ Alum Adjuvant through SP (subcutaneous injection). Blood was collected one week after the sixth immunization through heart puncture and clarified by centrifugation at 14,000 rpm for 30 min at 4 °C. Supernatant was aliquoted and stored at -80 °C. Where indicated this polyclonal serum was used at 500-fold dilution with 5% milk in TBST for primary antibody for western blot.

### Co-immunoprecipitation experiments and immunoblots

3T3-NTZ cells were transduced with retrovirus expressing mutants of FLAG tagged UNG and puromycin selected for 48 hours. Cells were lysed with lysis buffer (20 mM Tris-HCl (pH 7.5) 150 mM NaCl, 1 mM EDTA 1% NP-40) plus protease inhibitors with rocking in cold room for 30 min. Lysed cells were centrifuged twice at 4C with 15,000 rpm for 5min, supernatant were collected and quantified with BCA assay (Thermo Fisher Scientific, Catalog#: 23224, 23228). 1 mg of cell lysate were used to incubate with ANTI-FLAG® M2 Affinity Gel (Sigma Catalog # A2220-5ML) for 4 hours at 4 °C and beads were washed 5 times with lysis buffer. Beads were resuspended with 2X NuPAGE gel loading dye from Invitrogen and resolved on 12% NuPAGE along with 80 μg cell lysate as an input control. Proteins were transferred to 0.45 μM PVDF membrane (Thermo Fisher Scientific, Catalog#: 88518).and probed with anti-RPA32 antibody (4E4), then membrane were stripped and reprobed with anti-FLAG antibody (Sigma catalog # F7425-.2MG) for UNG detection. Anti-tubulin antibody was used as a loading control for lysate immunoblot after detection of RPA32 and UNG.

For co-immunoprecipitation of RPA mutants with FLAG-UNG. BOSC-23 cells were co-transfected with pMX-FLAG-mUNG and pMX-myc-RPA WT or pMX-RPA E252R, reciprocal IP was performed with anti-FLAG agarose followed by western blot of anti-RPA32 (4E4), stripped, reprobed with anti-FLAG, or IP performed with anti-myc plus protein G followed by western blot of anti-FLAG, stripped, reprobed with anti-RPA32. The NMR structure of human RPA winged helix domain in a complex with a peptide from RPA binding motif of human UNG (PDB 1DPU) was labeled and visualized with Pymol. The sites that were utilized in this study to disrupt the UNG-RPA interaction were highlighted as indicated. Mutation analysis was statistically tested by a two-tailed student T-test assuming unequal variance. P<0.05 considered to be significant.

## Acknowledgements

We thank Pamela Whitney for cell sorting. Dr. Xuesong Li from MDACC Cytogenetic and Cell Authentication Core for cell line authentication and mycoplasma check. Yunxiang Mu was supported by a fellowship from the UTMDACC Center for Cancer Epigenetics. National Institutes of Health [R01AI12539 to K.M.M.; R21 Al111129 to L.T.K and K.M.M.]. Recombinant Antibody Production Core [CPRIT RP190507]. Virginia Harris Cockrell endowment fund. Texas Tobacco Settlement – Molecular Mechanisms of Tobacco Carcinogenesis. This research was supported in part by the intramural Research Programs of the NIH, the National Cancer Institute (NCI) (L.T.K)

## Author Contributions

Y.M., L.T.K. and K.M.M. conceived the project. Y.M., Z.C., M.A.Z, J.B.P. performed experiments. Q.D. contributed reagents advice and critical intellectual input. Y.M. and K.M.M. assembled the figures and wrote the manuscript with input from all authors. L.T.K and K.M.M. acquired funding and supervised the work.

## SUPPLEMENTARY

**Table S1.**
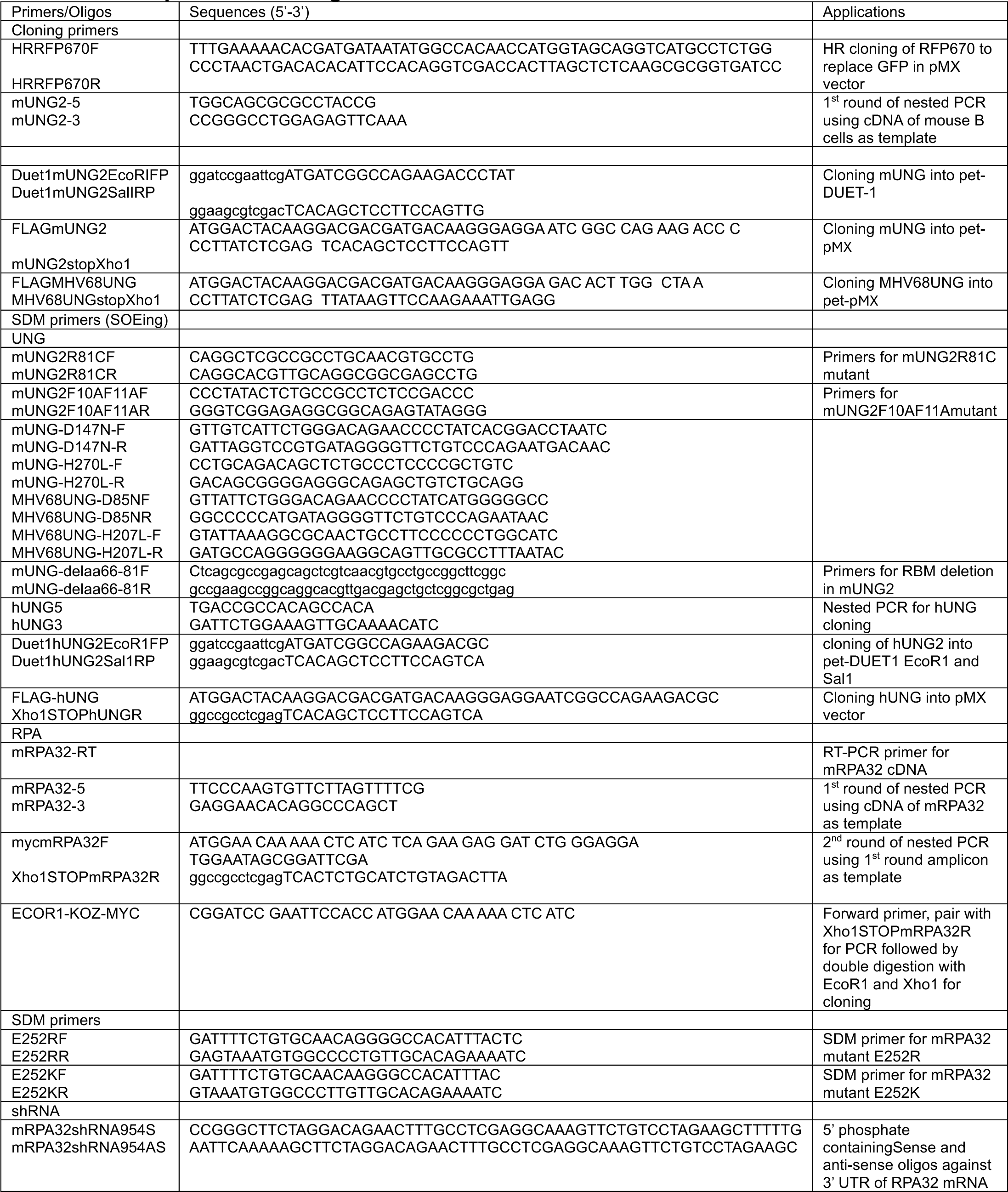
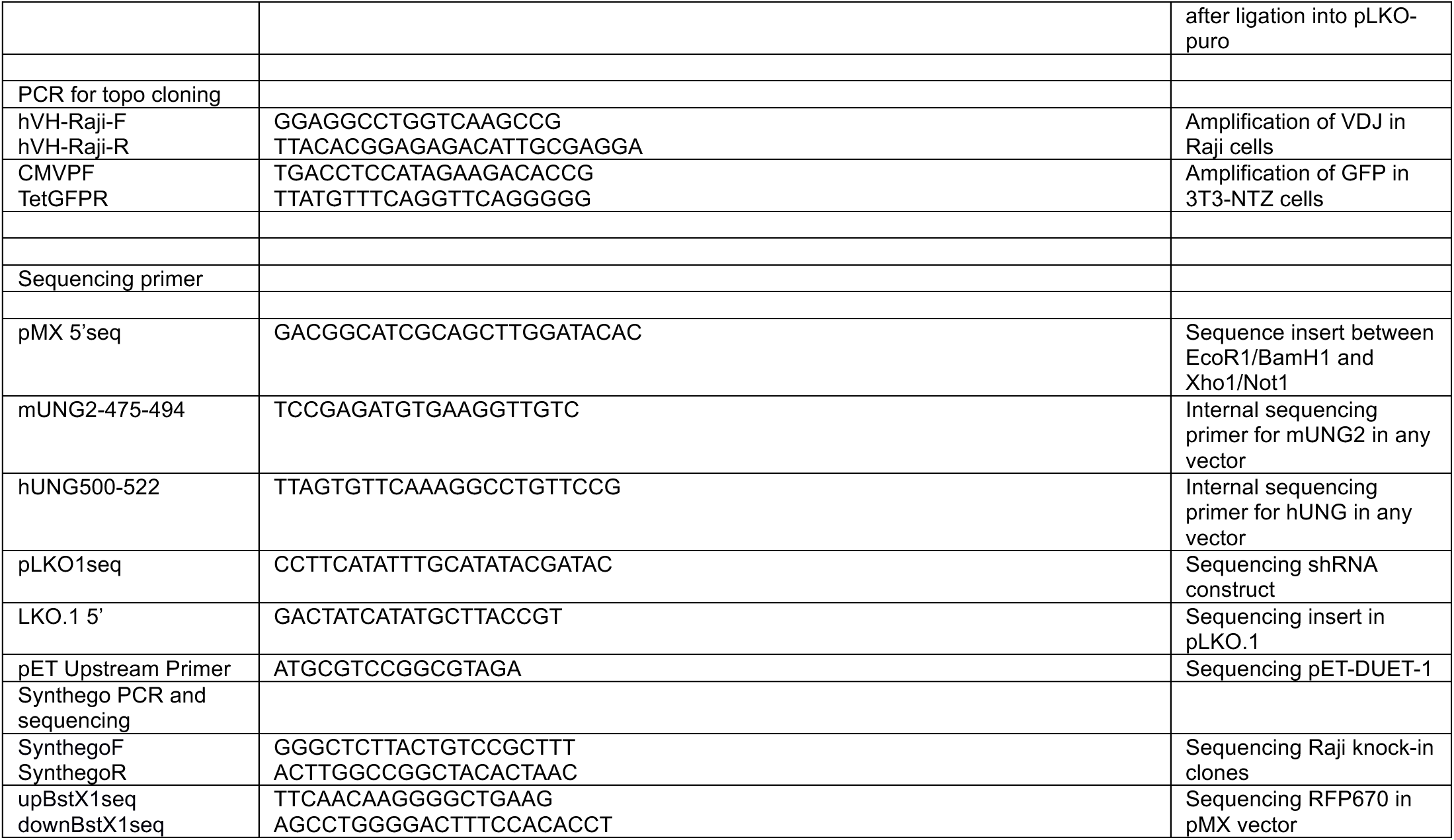
DNA primers and oligonucleotides.

**Figure S1.**
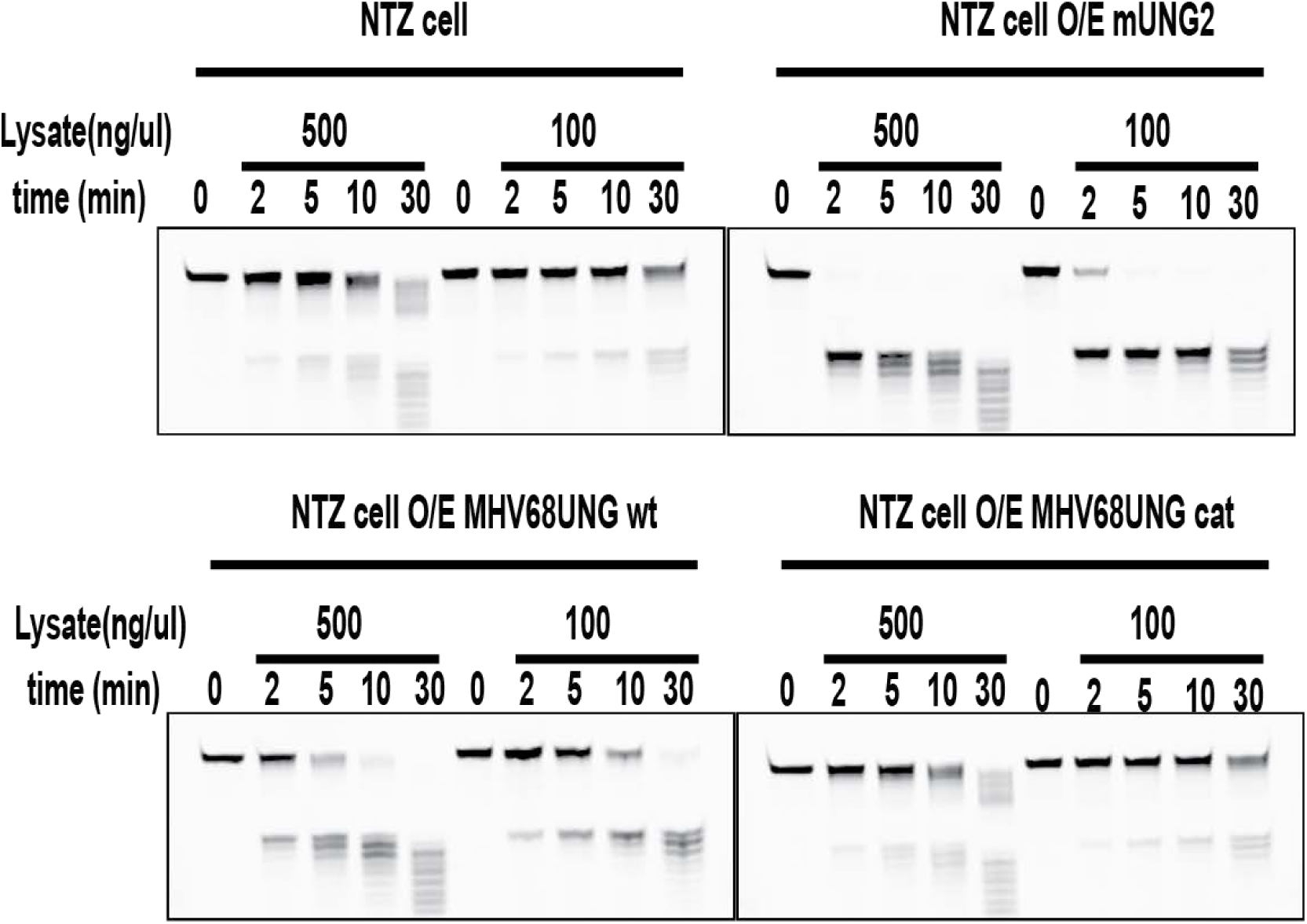
UDGase activity in cell lysates from 3T3-NTZ expressing UNGs. Lysates from 3T3-NTZ cells over expressing (O/E) the indicated UNGs, were evaluated in an oligo breakage assay. A G:U containing dsDNA oligo was incubated for the indicated time with 500 or 100 ng/ml lysates. UDGase activity was detected by conversion to faster migrating oligo after being resolved on 16% 8M Urea-PAGE gel.

**Figure S2.**
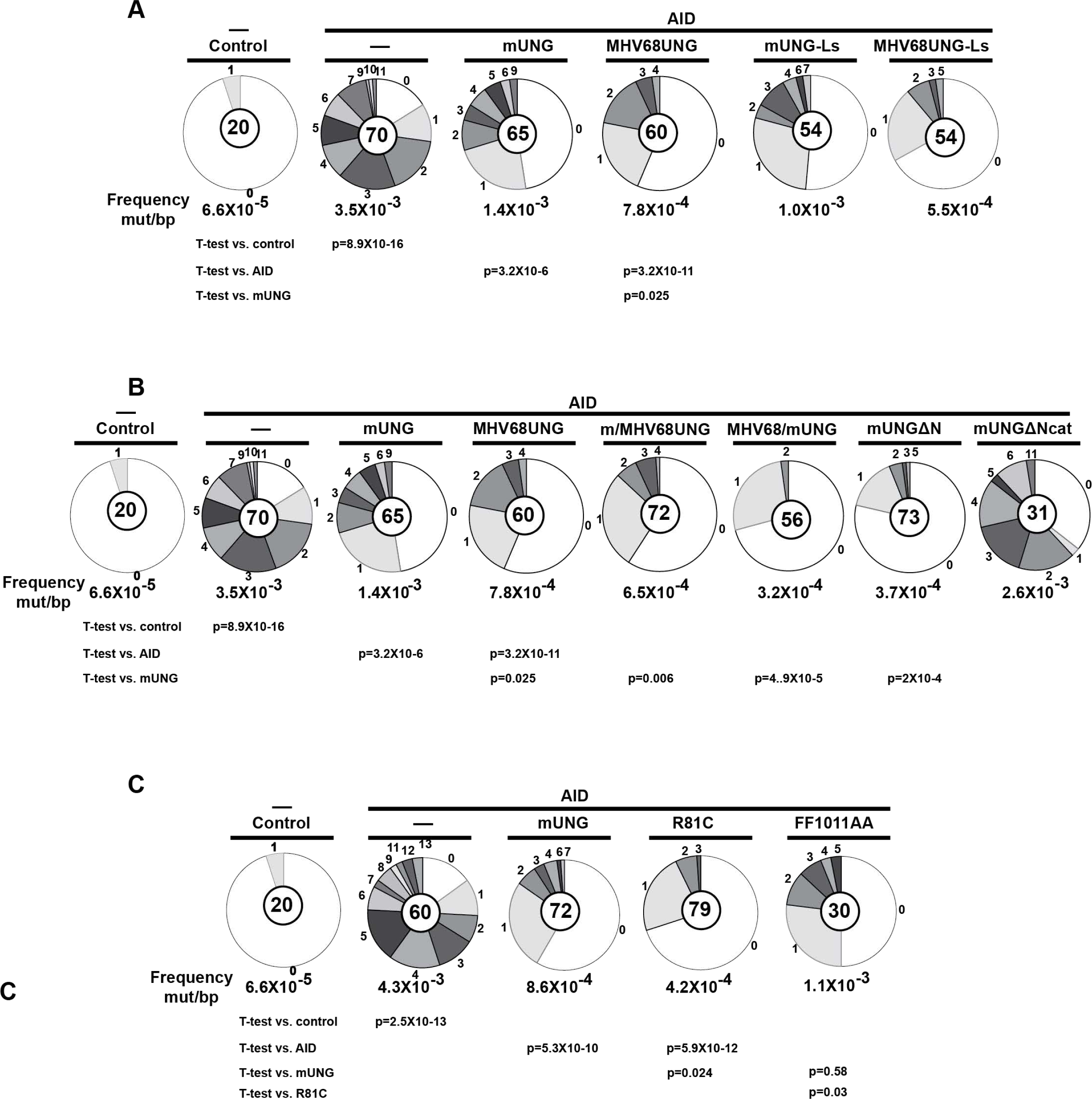
Mutation distribution and frequencies of MHV68UNG, mUNG and UNG mutants. Mutations in the GFP indicator gene in 3T3-NTZ mutation assay. Total mutation frequency is indicated below pie charts as mutations per bp. Segment size in pie chart is the proportion of clones with indicated number of mutations. The number of clones analyzed is indicated in the center. The presence of AID and which UNG is indicated. Control is indicative of background mutations likely from PCR error-rate. P-values (*, P < 0.05) were determined by a two-tailed t test assuming unequal variance. Displayed are mutations distribution and frequency from (A) Figure 3A: Comparative analysis of UNG leucine loop swap mutant. (B) Figure 3B: Analysis of UNG N-terminal mutants. Note: Data from A & B are from the same experimental sets and controls, mUNG and MHV68UNG are duplicated for comparative purpose in A & B. (C) Figure 4A: Analysis of mUNG R81C and mUNG F10A/F11AA mutants. Data was summary of n=1 (F10A/F11A) or n=3 (R81C) experiments as indicated in main figures.

**Figure S3.**
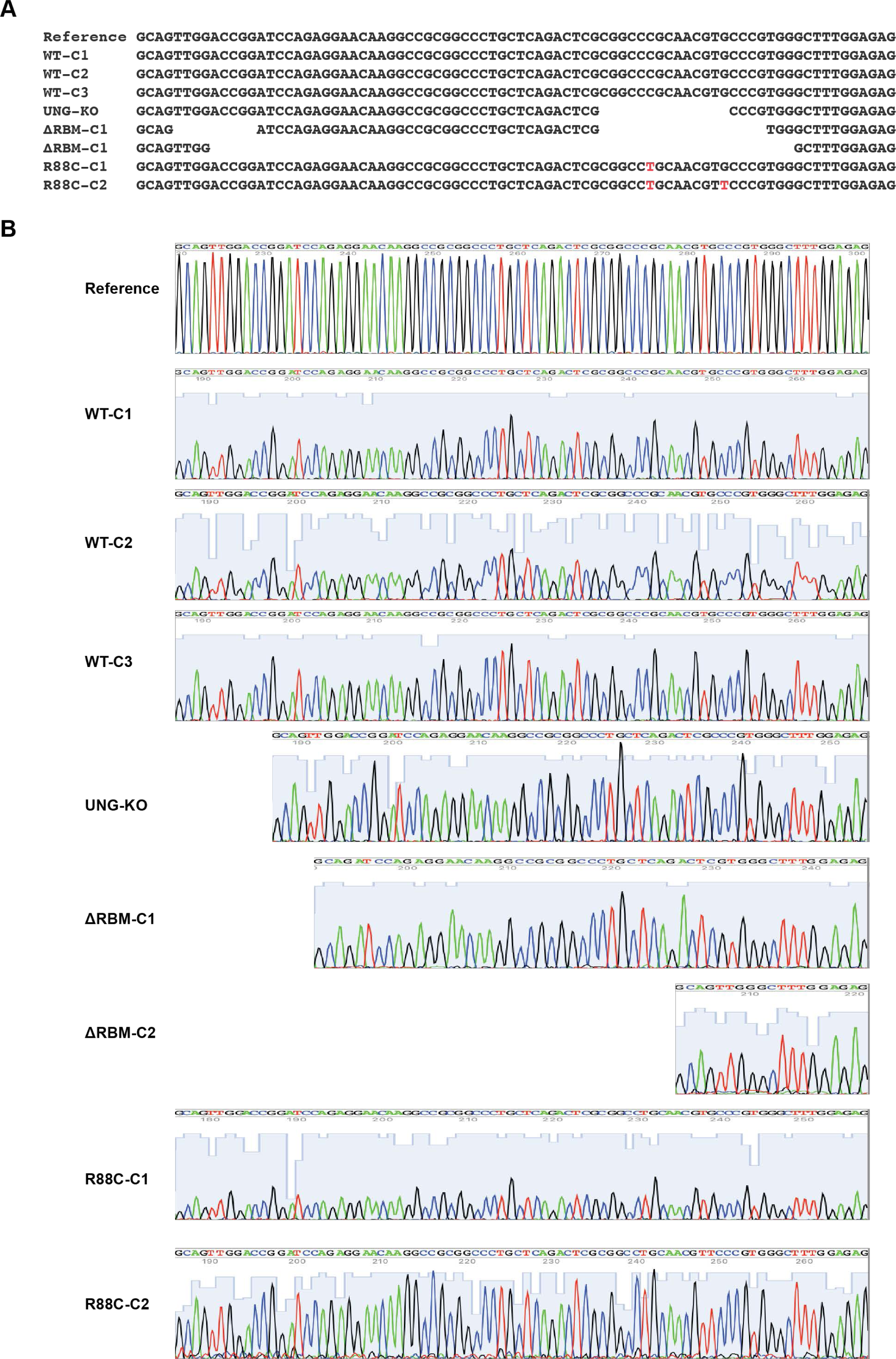
Raji single clones sequence confirmation of mutations in RBM domain. (A) Aligned nucleotide sequences of the Raji clones corresponding to the RBM coding region. Deletions (gaps) and mutations (red) are noted. The G to T mutation in R88C-C2 is a silent mutation. (B) Sequencing electropherograms of the RBM region from each clone is displayed.

**Figure S4.**
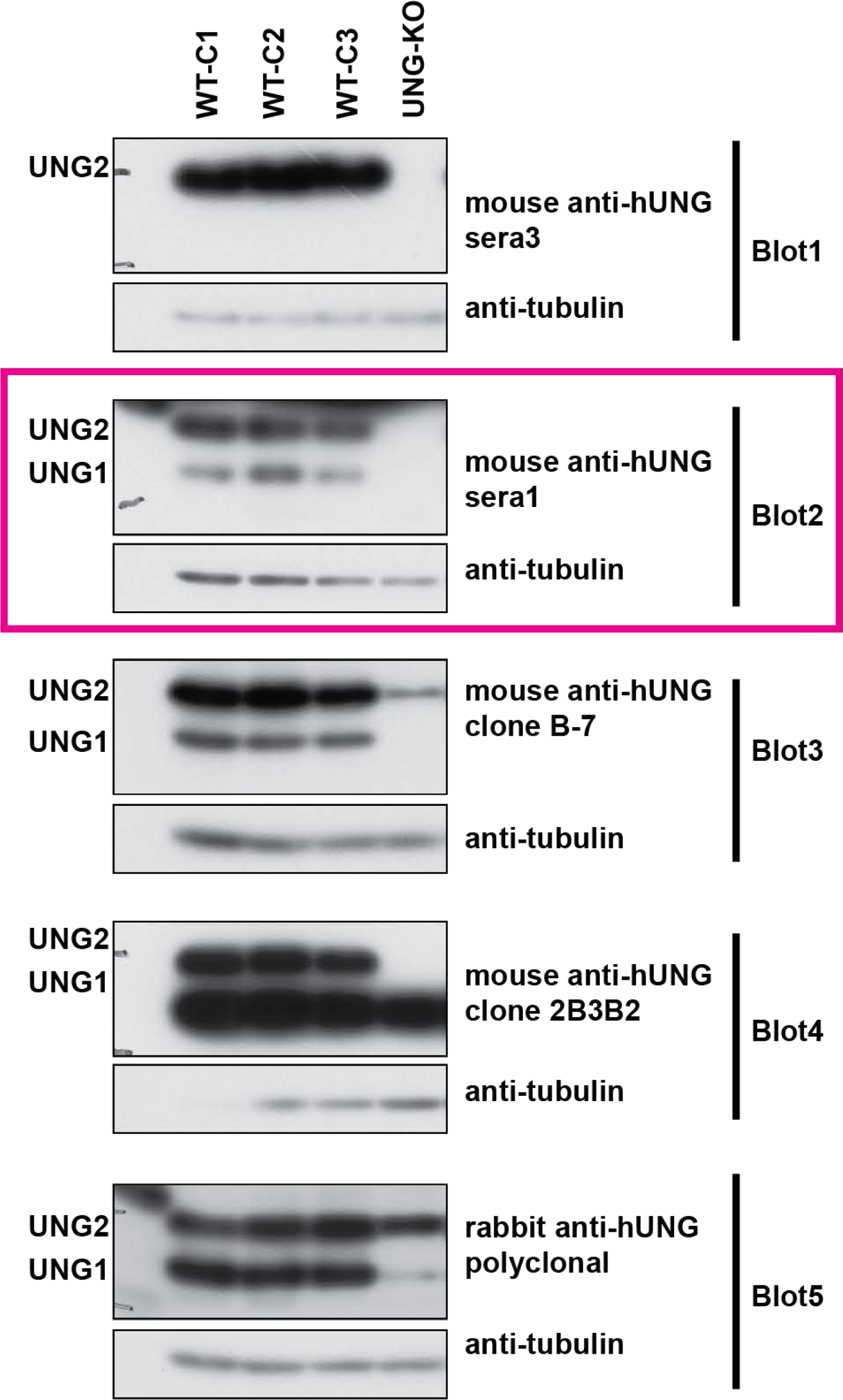
Immunoblot evaluation of UNG isoform detection by various antibodies. Lysates from the indicated Raji cell clones were immunoblotted with the indicated anti-hUNG antibody and anti-tubulin as a loading control. The migration of UNG1 (mitochondrial) and UNG2 (nuclear) isoforms are indicated. Only mouse polyclonal sera 1 (red box) clearly detected both isoforms without non-specific bands that impeded clear detection.

**Figure S5.**
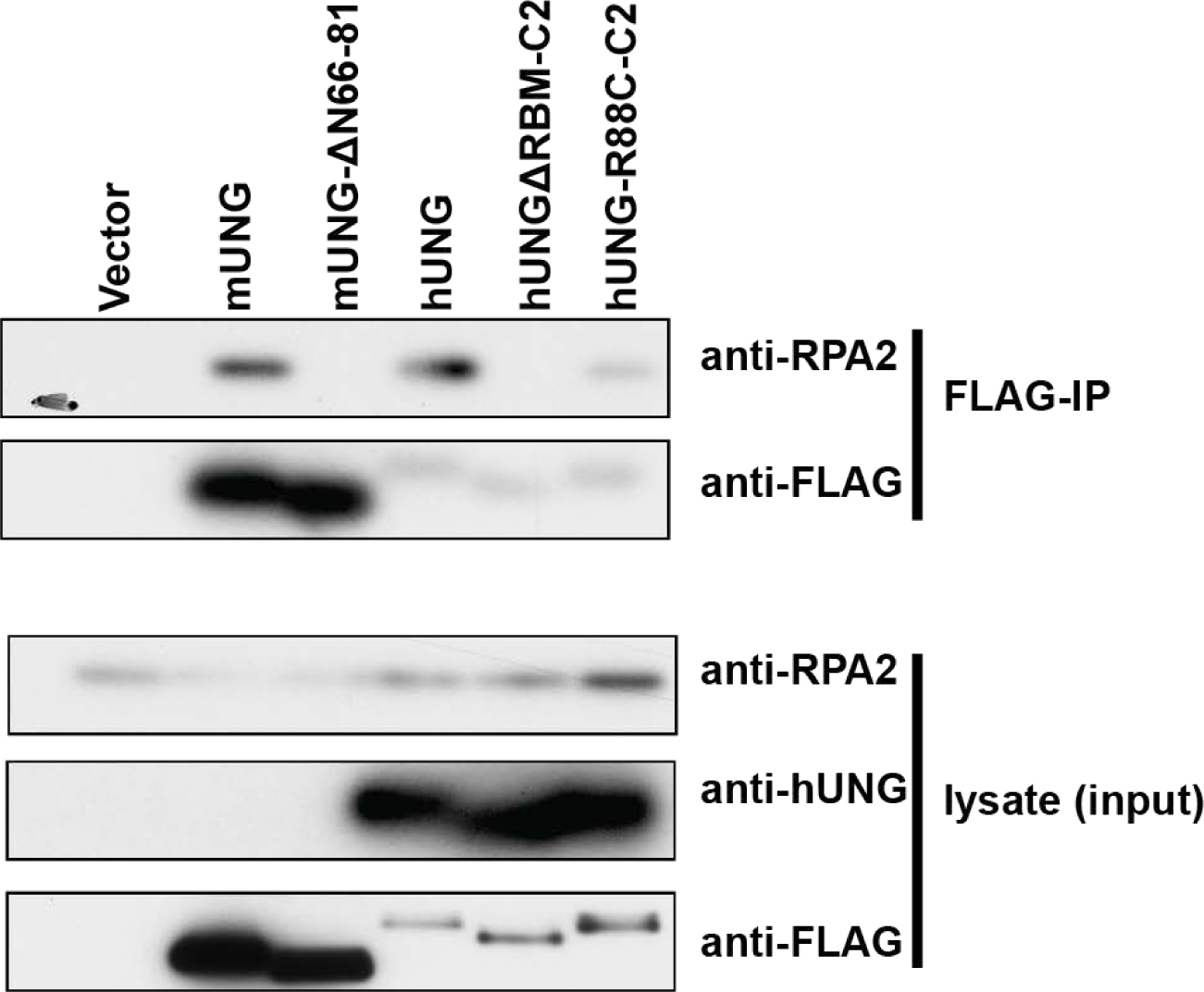
Degree of UNG interaction with endogenous RPA32 with hUNG mutants. Anti-FLAG co-IP from lysates of cells over-expressing indicated Flag-tagged UNGs. hUNGs were cloned from cDNA of Raji clones and transfected in human BOSC cells. UNGs were detected with anti-Flag and endogenous RPA32 was detected in anti-RPA immunoblot. mUNG ΔRBM66-81 has the region deleted that corresponds to the deletion in hUNG ΔRBM-C2.

